# Neuron-Specific WWOX Gene Therapy Produces Dose-Dependent, Durable Rescue in a Model of WWOX-Related Epileptic Encephalopathy

**DOI:** 10.64898/2026.03.11.710995

**Authors:** Mustafa Obeid, Rania Akkawi, Srinivasarao Repudi, Prince Kumar Singh, Baraa Abudiab, Allyson Berent, Thomas Brennan, Yael Weiss, Tawfeeq Shekh-Ahmad, Rami I. Aqeilan

**Affiliations:** The Concern Foundation Laboratories, The Lautenberg Center for Immunology and Cancer Research, Department of Immunology and Cancer Research-IMRIC, Hebrew University-Hadassah Medical School, Jerusalem, Israel; Institute for Stem Cell Science and Regenerative Medicine, Bangalore 560065 Karnataka, India; Institute for Drug Research, The School of Pharmacy, Faculty of Medicine, The Hebrew University of Jerusalem, Jerusalem 91120, Israel; Mahzi Therapeutics, South San Francisco, United States

**Keywords:** Gene therapy, WOREE syndrome, DEE, WWOX, WPRE

## Abstract

Biallelic loss-of-function mutations in WWOX cause a spectrum of neurodevelopmental disorders, including the severe, early-onset WOREE syndrome, frequently associated with intractable epilepsy and premature mortality, and the milder SCAR12, characterized by subtler neurological manifestations. While neuronal replacement of WWOX has emerged as a potential therapeutic strategy, the parameters required for safe, durable, and clinically translatable gene delivery remain undefined. Here, we systematically delineate the determinants of effective WWOX gene therapy by evaluating promoter selection, cellular targeting, vector configuration, dose, and developmental timing in a severe *Wwox*-null mouse model. Neuron-restricted expression driven by the human Synapsin I promoter uniquely enabled sustained phenotypic correction, whereas ubiquitous or oligodendrocyte-restricted expression failed to confer durable benefit. To better regulate transgene expression, we generated a vector lacking the WPRE element, enabling dose calibration within a clinically relevant range. Comparative dose-response analyses identified an optimal therapeutic dose of AAV9-hSynI-WWOX that produced robust, dose-dependent rescue of survival and growth. Moreover, the rescued mice displayed glucose, behavioral function and fertility indistinguishable from WT mice, accompanied by long-term restoration of WWOX DNA, transcript, and protein across central and peripheral neural tissues without detectable hepatic expression. Neuronal WWOX reconstitution promoted widespread myelination and attenuated neuroinflammatory responses, including astrogliosis and microglial activation to levels indistinguishable from WT. Continuous electrocorticographic monitoring uncovered early postnatal neuronal hyperexcitability in *Wwox*-null mice, which was effectively suppressed by therapeutic WWOX delivery. Finally, our data define an early postnatal therapeutic window in *Wwox*-null mice, showing that treatment initiated between postnatal days 1 and 5 supports durable rescue. Together, these findings define a rigorously optimized, neuron-targeted AAV9-WWOX gene therapy framework and establish critical design and timing principles for translational treatment of WWOX-associated developmental and epileptic encephalopathies.

## Introduction

The WW domain-containing oxidoreductase (*WWOX*) gene spans one of the most active common fragile sites. Common fragile sites have been suggested to play roles in a wide range of neurodevelopmental processes, including cell-cell adhesion, synapse organization, interneuron differentiation, neuronal projection development, glutamate receptor signaling, and CNS axonogenesis ^1^. *WWOX* maps to chromosome 16q23.3-24.1 and encodes a 46-kDa protein composed of 414 amino acids, with key roles in tumor suppression, DNA damage response, and metabolic regulation^2,3^. Structurally, WWOX features two N-terminal WW domains, a nuclear localization signal (NLS), and a C-terminal short-chain dehydrogenase/reductase (SDR) domain, enabling it to interact with multiple protein partners and engage in diverse signaling pathways^4–7^.

*WWOX* has been extensively studied for its tumor suppressor functions, primarily through regulation of key cancer-associated signaling pathways, including WNT, TGFβ1, NF-κB, and receptor tyrosine kinases (RTKs)^8–12^. In addition, *WWOX* modulates cellular metabolism via HIF1α and IDH pathways^13,14^, and contributes to the DNA damage response through interactions with ATM^14^, BRCA1^15,16^, and p73^17,18^. These activities highlight its key role in tumor formation. More recently, increasing evidence indicates a vital role for WWOX in the nervous system. WWOX is highly expressed in the brain, showing dynamic patterns during embryonic and postnatal development, and is crucial for maintaining the central nervous system (CNS) homeostasis^8,19–23^.

Mutations in *WWOX* have been linked to a broad spectrum of neurological disorders, including Alzheimer’s disease^24–26^, Parkinson’s disease^27,28^, multiple sclerosis^29–31^, autism spectrum disorders^32,33^, epilepsy, and ataxia^34^, underscoring its essential role in CNS health and function. Of particular interest, germline recessive mutations including missense, nonsense, and deletion variants are causally associated with two major inherited syndromes: SCAR12 (spinocerebellar ataxia, autosomal recessive 12; OMIM 614322) and WOREE syndrome (WWOX-related epileptic encephalopathy, also classified as developmental and epileptic encephalopathy 28; OMIM 616211).

WOREE syndrome is a severe, early-onset neurological disorder caused primarily by biallelic mutations that result in premature stop codons or complete loss of *WWOX* expression. Clinical features include profound global developmental delay and intractable epilepsy, with seizure types such as tonic, clonic, tonic-clonic, myoclonic, infantile spasms, and absence seizures. Most affected individuals fail to reach basic developmental milestones, including eye contact, sitting, speech, or ambulation. Approximately 50% of WOREE patients harboring null mutations die by 5 years of age. In contrast, SCAR12 presents with a milder phenotype, often involving early-onset epilepsy and cerebellar ataxia that may respond to anti-epileptic drugs (AEDs), although patients frequently experience intellectual disability and motor dysfunction. *WWOX* mutations have also been implicated in West syndrome, which is characterized by infantile spasms and hypsarrhythmia. Magnetic resonance imaging (MRI) of children with *WWOX* mutations commonly reveals structural abnormalities such as corpus callosum hypoplasia, delayed myelination, progressive cerebral atrophy, and optic nerve atrophy^35–41^.

Previous work from our laboratory sought to define the cell populations underlying the WWOX-deficient phenotype. Using conditional mouse models, we showed that deletion of *Wwox* in neural stem and progenitor cells driven by the Nestin promoter (N-KO), or in postmitotic neurons driven by the Synapsin promoter (S-KO), leads to severe epilepsy, hypomyelination, ataxia, and early lethality by 3-4 weeks of age, closely phenocopying global *Wwox*-null mice^42^. These observations established neuronal WWOX as indispensable for normal brain function and survival and provided the rationale for testing whether targeted restoration of WWOX expression could reverse these profound neurological deficits^43,44^.

Neuron-targeted AAV9-mediated delivery of WWOX achieves widespread neuronal expression in the developing brain and ameliorates disease manifestations in Wwox-deficient mice^45^. However, key gaps remained regarding optimal dosing and safety, prompting the present study. In this study, we refine the *WWOX* gene therapy approach to improve effectiveness for potential clinical use. By driving *WWOX* expression under the neuron-specific Synapsin I promoter, we achieve targeted and strong neuronal transduction. To better regulate transgene expression, we generated a vector lacking the WPRE element. Additionally, we demonstrated a dose-dependent therapeutic effect, with neuronal WWOX restoration in *Wwox*-null mice rescuing key phenotypes, including premature death, growth delay, hypoglycemia, and behavioral deficits to levels indistinguishable from WT mice. Furthermore, this approach significantly enhances CNS myelination, decreases neuroinflammation, and lessens gliosis. These findings, along with growing evidence linking *WWOX* mutations to various neurological disorders, highlight the potential of *WWOX* gene replacement therapy. Overall, our results provide a solid foundation for developing effective treatments for WOREE syndrome, SCAR12, and likely other *WWOX*-related neurodevelopmental and neurodegenerative diseases.

## Materials and methods

### Plasmid vectors

Human WWOX cDNA was cloned under different promoters, including human Synapsin I (hSynI), EF1-alpha, CBA, MBP, and CMV, in the pAAV. These vectors were packaged into AAV9 serotypes (Fujifilm Diosynth Biotechnologies, Watertown, Massachusetts, USA), (Vector Biolabs, Philadelphia, USA). Custom-made AAV9-CBA-hWWOX and AAV9-hSynI-EGFP viral particles were obtained from the Vector ELSC Core Facility at the Hebrew University of Jerusalem. The viral titer was measured by qRT-PCR using bGH primers.

### Mice

The generation of *Wwox*-null ^(−/−)^ mice (KO) was previously reported^46^. Mice were kept on an FVB background. Heterozygote ^(+/−)^ or KO rescued mice (KO injected with AAV9-hSynI-hWWOX) were used for breeding to generate KO mice. After weaning, wild-type (WT) and KO rescued mice were separated by sex and housed in individual ventilated cages (IVC). To assess fertility, one male and one female of the same genotype were housed together in an IVC cage. All animals were maintained in a specific pathogen-free (SPF) facility under a 12-hour light/dark cycle, with *ad libitum* access to food and water. All animal experiments were conducted in accordance with the ethical guidelines and with prior approval from the Hebrew University Institutional Animal Care and Use Committee (HU-IACUC).

### Genotyping

Genomic DNA was extracted from the tail/ear using DNA lysis buffer (20mM NaOH, 0.1mMEDTA). PCR reaction was performed using Taq DNA polymerase (2x rapid Taq master mix-P222-01-Vazyme) and the following primers were used: shared 5’ primer of the WT *Wwox* allele (WF, 5’ GCAGAATGTCTTGCTAGAGCTTTG 3’), 3’ primer in sequence deleted in the targeted allele (WR, 5’ ATACTGACATCTGCCTCTAC 3’), and a 3’ primer internal to the Geo cassette (GeoR1, 5’ CAAAAGGGTCTTTGAGCACCAGAG 3’).

### Intracerebroventricular (ICV) injection of AAV particles into P0-P5 *Wwox*-null mice

Intracerebroventricular injections of either AAV9-hWWOX (under different promoters) or reference item (RI) were conducted using stereotactic technique to ensure consistency following the published protocol^47^. Briefly, *Wwox*-null (KO) or WT neonates were anesthetized by placing them on a dry, flat, cold surface. The anesthetized pup head was gently wiped with a cotton swab soaked in 70% ethanol. The pup (KO or WT) was staged on the stereotaxic frame for AAV9-hWWOX or RI delivery. The injection site was targeted at x = ±0.8 mm, y = 1.5 mm, and z = -1.6 mm, relative to lambda. These anatomical landmarks were visible through the skin at P0-P1. For older pups, the coordinates were adjusted to account for developmental changes; for P5-pups , x = ±1.0 mm, y = 1.0 mm, and z = -2.0 mm. A Micro-4 nano-pump controller was used to ensure a steady injection rate of 1-1.5 μl/min, delivering 2.0 μl/hemisphere using a Hamilton syringe with a 32G needle (World Precision Instruments). The needle was kept at the injection site for 30-60 seconds to allow proper diffusion and then removed slowly over 1 minute. The procedure was repeated for the contralateral hemisphere. After the injection, the pups were placed on a warming pad till they awoke and then returned to their mother’s cage. The pups were closely monitored for growth rate, glucose levels, seizures, ataxia, and general condition to assess phenotypes.

### Blood glucose levels

Blood glucose was measured at p10, p20, and p30 from the tail nick using a glucometer (Roche Diagnostics, Mannheim, Germany).

### DNA and RNA extraction

Brain tissue samples were collected, snap-frozen in liquid nitrogen and preserved in -80℃ for analysis. DNA extraction was performed using the DNeasy Blood and Tissue Kit (Qiagen) according to the manufacturer’s protocol. Briefly, tissue samples (up to 25 mg) were lysed in ATL buffer supplemented with Proteinase K at 56°C overnight. Following lysis, samples were mixed with AL buffer and 100% ethanol to facilitate DNA binding to the DNeasy Mini Spin Column. The bound DNA was washed sequentially with AW1 and AW2 buffers to remove contaminants and ensure that only pure DNA remains bound to the column. Then AE buffer was added to the mini spin column and incubated for 5 minutes at RT for elution of purified DNA. DNA integrity and concentration were assessed prior to downstream applications using DeNovix (DS-11FX+). For qPCR, 50 ng of DNA template was used per reaction. The primer sequences were as follows: forward 5′-GCTCTCTTAAGGTAGCCCCG-3′, reverse 5′-CGCCTCATCCTGGTCCTAAA-3′.

Total RNA was isolated from non-perfused tissue using TRIzol reagent (Bio-Tri) according to the manufacturer’s phenol-chloroform protocol. Briefly, tissues were homogenized in TRIzol at 1 mL per 50-100 mg tissue, followed by phase separation with chloroform. The aqueous phase was recovered, RNA was precipitated with isopropanol, the pellet was washed with 75% ethanol, air-dried, and resuspended in DEPC-treated nuclease-free water. RNA concentration and integrity were assessed on a DeNovix (S-11 FX+) spectrophotometer. Extracted RNA was stored at -80 °C until use. cDNA was synthesized from 1 µg total RNA using the qScript cDNA Synthesis Kit (Quantabio) following the manufacturer’s instructions. For qPCR, 50 ng of cDNA template was used per reaction. The primer sequences were as follows:

forward, 5′-ACCTACTTGGACCCAAGACTGGCG-3′;
reverse, 5′-GGTGCTGCCGTCGTATCTTTGCC-3′.

### Immunoblot

Brain tissue samples were homogenized using protein lysis buffer (50mM Tris, PH 7.5, 150mM NaCL, 10% glycerol, 0.5% Nonidet P-40 (NP-40) with the addition of protease and phosphatase inhibitors. The lysates were subjected to SDS-PAGE under standard conditions. The following antibodies were used: rabbit polyclonal anti-WWOX 1:10,000, mouse monoclonal anti-GAPDH (CB-1001), Calbiochem 1:10,000, rabbit polyclonal anti-HSP90 (4874S), cell signaling 1:1000.

### Immunohistochemistry

Mice (WT, KO, KO injected mice) at different ages (p10-p180) were deeply anesthetized using an overdose of ketamine and xylazine and then transcardially perfused using 4% PFA/PBS. Dissected brains were post-fixed in 4% PFA overnight at 4°C, followed by incubation in 30% sucrose overnight at 4°C, and were then embedded in O.C.T. Sagittal brain sections (14 μm) were prepared using a cryostat and can be stored at -80°C. Sections were warmed to room temperature, washed with PBS and fixed with 4% PFA for 5 min, followed by PBS washing. Permeabilization was done using PBT (0.1% Triton X-100 in PBS). Formalin-fixed, paraffin-embedded (FFPE) sagittal tissue sections (14um) were deparaffinized, followed by antigen retrieval with 25 mM citrate buffer (pH=6) in a pressure cooker at high pressure for 3 minutes. The sections were then left to cool for 25 minutes. Fresh coronal sections were prepared using a vibratome (50 µm). The sections (FFPE, frozen, fresh sections) were then washed and blocked using a blocking buffer (5% normal goat serum, 0.5% BSA and 0.1 %Triton X-100 in PBT) followed by incubation with a specific primary antibody in a humidified chamber overnight (rabbit polyclonal anti-WWOX 1:5,000, mouse anti-beta III tubulin (B364854), biolegend,1:1000), rabbit anti-GFAP-clone EPR1034Y (AB68428), Abcam 1:250, rabbit anti-Iba1-clone HL22 (ab28937), Abcam 1:250, rabbit anti-MBP, (Ab65988), Abcam 1:200, mouse anti-NeuN, (MAB377), Milipore, 1:500). The sections were washed using PBST buffer (0.05% Tween-20 in PBS) and incubated with the corresponding secondary fluorescent antibody for 1hr at room temperature (goat anti-rabbit Alexa flour 647 (Ab150079), Abcam 1:1000) and goat anti-mouse Alexa flour 488 (ab150117), Abcam 1:1000) with Hoechst 33258 (1:1000). Then the sections were washed with PBST three times and were mounted using fluorescent mounting media (Dako s3023). Sections were then imaged using a 3DHISTECH panoramic scanner (3DHISTECH, Hungary).

### Sciatic Nerve and spinal cord Extraction

Sciatic nerve extraction was performed following the previously published protocol^48^. Briefly, in a transcardially perfused mouse, a dorsal incision along the hind leg is made to expose the sciatic nerve. Using fine-point tweezers and surgical scissors, the sciatic nerve is carefully dissected from its surrounding tissues. The nerve is handled delicately to avoid damage. For the spinal cord extraction, a midline incision along the dorsal part of the mouse was made with a scalpel. Then, the skin, underlying muscles, and bones were carefully removed to expose the spinal cord. Using forceps, the spinal cord was gently extracted from the vertebral column. Immediately after excision, the specimen (sciatic nerve, spinal cord) is either snap frozen for further analysis or immersed in 4% formalin to ensure rapid and uniform fixation for 24-48 hours at RT. After fixation, the tissue was transferred to 70% ethanol for 24 hours prior to processing in an automated tissue processor (Leica, Histocore). Then, the tissue was embedded in paraffin blocks, and sections were prepared using a microtome (14 μm).

### Behavioral tests

Behavioral assessment was conducted on WT+RI and KO rescued mice (HD) at the age of 3-months (both males and females) as previously published^45^. Animals were assessed in an open-field test, an elevated plus maze, and a rotarod. The open-field test was conducted following a previously published protocol^49^. Briefly, mice were placed in a 50 × 50 × 33 cm arena and allowed to explore freely for 6 minutes. The center of the arena was defined as a 25 × 25 cm square. Key measurements, including the mice velocity, time spent in the central area, and time spent near the arena’s perimeter, were recorded. The test was monitored using a video camera connected to a computer with tracking software (Ethovision 12). The elevated plus maze consisted of two open arms (30 × 5 cm) with a 1 cm high rim and two closed arms (30 × 5 cm) with a 16 cm high rim, positioned perpendicular to each other. The maze was elevated 75 cm above the floor.

Mice were placed into the maze and allowed to explore freely for 6 minutes. The time spent in both the open and closed arms was recorded to assess anxiety-like behaviors. For motor coordination and balance assessment, a rotarod test was performed. Each mouse was placed on a rotating rod, which gradually accelerated from 5 revolutions per minute (rpm) to 40 rpm over 99 seconds. Mice underwent three trials, with a 20-minute interval between trials. The latency to fall off the rod was recorded, and if the mouse remained on the rod for 240 seconds without falling, the trial was terminated. Results from all trials were used to calculate the average latency to fall.

### DRG culture

Neurons were cultured following a previously published protocol^45^. Briefly, DRG neurons were isolated from mouse embryos at E13.5. Embryos were genotyped and the DRGs were collected in cold L-15 medium. Tissues were dissociated in 0.25% trypsin, triturated, centrifuged, and re-suspended in NB medium (Neurobasal, B27 supplement, 0.5 mM l-glutamine, and penicillin–streptomycin). Pre-cleaned 13-mm diameter glass coverslips were placed in 4-well dishes and coated with Matrigel (1 h at RT) then poly-D-lysine (30 min at RT) prior to dissection. Cells were plated at a density of 40,000 cells/13-mm coverslips in NB medium and maintained in a humidified incubator at 37°C and 5% CO2. Cultures were treated with fluorodeoxyuridine at (Days In Vitro) DIV2, 4, and 6 to eliminate non-neuronal cells. Fifty percent of the cell culture media was replaced every third day. At DIV6, *Wwox*-null DRG neurons were infected with AAV9-SynI-EGFP or AAV9-SynI/CMV/MBP-hWWOX and cultured for 3 days, and then, expression of GFP and WWOX were validated by immunohistochemistry.

### Surgery and ECoG data acquisition

This study utilized WT, WWOX knockout (KO) mice injected with RI, and WWOX-KO mice injected with AAV9-hSynI-hWWOX (rescue group). At P14, pups were anesthetized with isoflurane (1–2%, adjusted for low body weight; Terrell™, USP, Piramal Critical Care) and positioned in a stereotaxic frame (Kopf Instruments, CA, USA). Carprofen was administered prior to surgery, and animals were implanted with a single-channel electrocorticography (ECoG) transmitter (model A3049F2; Open-Source Instruments, USA). Unlike standard adult procedures, the transmitter body was kept outside of the body to accommodate the small size of pups. Two intracranial electrodes were placed epidurally: the recording electrode was positioned above the right dorsal cortex, and the reference electrode was placed contralaterally. Electrodes were secured to the skull with miniature screws and tissue adhesive, and the incision was sealed with dental cement. Immediately after surgery, continuous wireless ECoG acquisition was initiated without a recovery interval. Recordings were maintained for 7 consecutive days, during which pups were housed with their mother and monitored under continuous 24/7 ECoG activity. Signals were acquired and stored using Neuroarchiver software (Open-Source Instruments Inc.). ECoG traces were manually screened for interictal spikes and spike–wave discharges (SWDs) by a blinded investigator. Events were defined based on amplitude, duration, and morphology criteria established in previous mouse epilepsy models. A subset of events was cross-validated with time-synchronized video recordings to confirm electrographic abnormalities.

### Statistics

Statistical analyses were performed using GraphPad Prism (version 5 or later) and Microsoft Excel. Data were analyzed in a blinded manner when feasible and are presented as mean ± standard deviation (SD), as indicated in the figure legends. Comparisons between two groups were performed using two-tailed unpaired Student’s t-tests. Survival was analyzed using Kaplan–Meier curves with significance assessed by the log-rank (Mantel–Cox) test. Longitudinal measurements (e.g., body weight and blood glucose) were analyzed at individual time points as specified in the figures. Electrocorticography (ECoG), behavioral, molecular, and histological quantifications were performed on per-animal averaged values and compared using unpaired t-tests. Exact sample sizes (n) and P values are reported in the figure legends. A P value < 0.05 was considered statistically significant; non-significant results are denoted as ns.

## Results

### Comparative promoter analysis identifies neuron-specific WWOX expression via the Synapsin promoter as optimal for disease rescue

Given the essential role of neuronal WWOX in CNS homeostasis, we systematically evaluated AAV-mediated human WWOX (hWWOX) expression using promoters selected to test expression breadth, strength, and cell-type specificity. Ubiquitous promoters (EF1α, CMV, CBA) were used to assess whether broad or strong expression could compensate for WWOX loss, whereas the MBP promoter tested oligodendrocyte-restricted rescue, and the Synapsin I promoter enabled neuron-specific restoration. Therapeutic efficacy was evaluated by survival, body weight, and blood glucose levels.

EF1α, a broadly active but relatively weak promoter^50^, drove widespread WWOX expression but failed to rescue lethality, growth retardation, or hypoglycemia in *Wwox*-null mice (Figure 1A-C,M, Figure S1A,B, Figure S2D). Stronger ubiquitous promoters (CMV and CBA)^51,52^, supported early postnatal rescue of survival and glycemic defects, indicating that early high-level expression is sufficient for transient benefit, but this protection was not sustained at later stages (Figure 1D-F,M, Figure S1A,B, Figure S2A-D). In contrast, oligodendrocyte-specific WWOX expression driven by the MBP promoter resulted in weak expression and no phenotypic improvement, demonstrating that restoration in this lineage alone is insufficient (Figure 1G-I,M, Figure S1A,B, Figure S2D).

**Figure 1:**
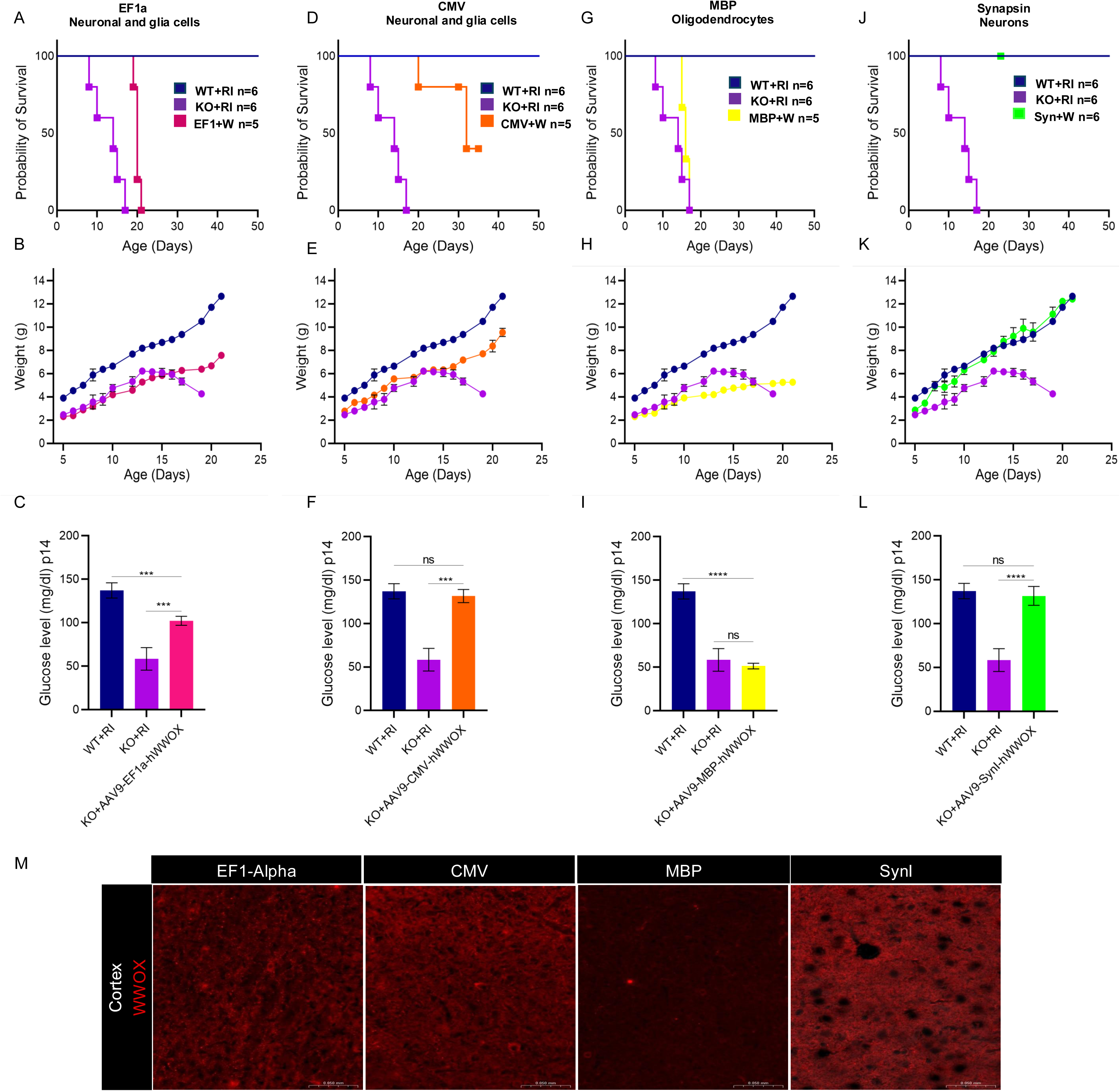
*WWOX* restoration driven by the Synapsin promoter achieves high expression efficiency. **(A,D,G,J)** Kaplan–Meier survival analyses of *Wwox*-null mice treated with AAV9-EF1a-hWWOX (4×10^10^ vg) (n = 5), AAV9-CMV-hWWOX (4×10^10^ vg) (n = 5), AAV9-MBP-hWWOX (4×10^10^ vg) (n = 5) and AAV9-hSynI-hWWOX (4×10^10^ vg) (n = 6), respectively compared to *Wwox*-null (KO) and wild type (WT) mice injected with the reference item (RI) (n = 6). p < 0.01, log-rank (Mantel–Cox) test. **(B,E,H,K)** Graphs showing body weight (g) of WT (n = 6), KO mice injected with the reference item (RI) (n = 6), and KO mice injected with hWWOX driven by different promoters (n=5-6) at multiple postnatal time points. At postnatal day 14 (P14), KO mice treated with hWWOX-hSynI and hWWOX-CMV exhibited significant increases in body weight compared with RI-injected KO mice (p < 0.001 and p < 0.01, respectively), whereas no statistically significant differences were observed in KO mice treated with hWWOX-EF1α or hWWOX-MBP. **(C,F,I,L)** Bar graphs representing glucose levels (mg/dl) at p14 in WT (n=6), KO injected with RI (n=6) and KO injected with hWWOX using different promoters (n=5-6). p<0.001 for KO injected with hWWOX-hSynI and hWWOX-CMV, but no significant difference was observed for KO injected with hWWOX-Ef1a and KO injected with hWWOX-MBP compared to KO injected with RI. **(M)** Immunohistochemistry staining for WWOX (red) in the cortex of p18 mice indicating the specificity and intensity of WWOX restoration using different promoters.

Strikingly, neuron-specific expression under the human Synapsin I promoter yielded the most robust and sustained rescue, with efficient neuronal expression in vivo (Figure 1J-M, Figure S1A-D, Figure S2D) and superior neuron-restricted transduction in primary neuronal cultures compared with CMV, while MBP-driven vectors showed no neuronal expression (Figure S2E). These findings align with our earlier conditional knockout models ^42^ and identify neuronal WWOX restoration via the Synapsin I promoter as the optimal and disease-relevant gene therapy strategy, which was therefore used for all subsequent experiments.

### Removal of WPRE enables controlled, physiologic WWOX expression

In our earlier therapeutic construct, we incorporated the woodchuck hepatitis virus post-transcriptional regulatory element (WPRE), a widely used enhancer of transgene expression that increases mRNA stability, promotes nuclear export, and improves transcript termination, typically resulting in a 5-8 fold increase in expression^53^. As expected, the presence of WPRE downstream of the transgene markedly augmented WWOX expression when compared to wild-type expression (Figure 2). To directly evaluate the intrinsic therapeutic efficacy of WWOX expression and enable tighter control of transgene output, we subsequently removed the WPRE element and assessed WWOX expression and the ability of WWOX alone to rescue phenotypes in *Wwox*-null mice.

**Figure 2:**
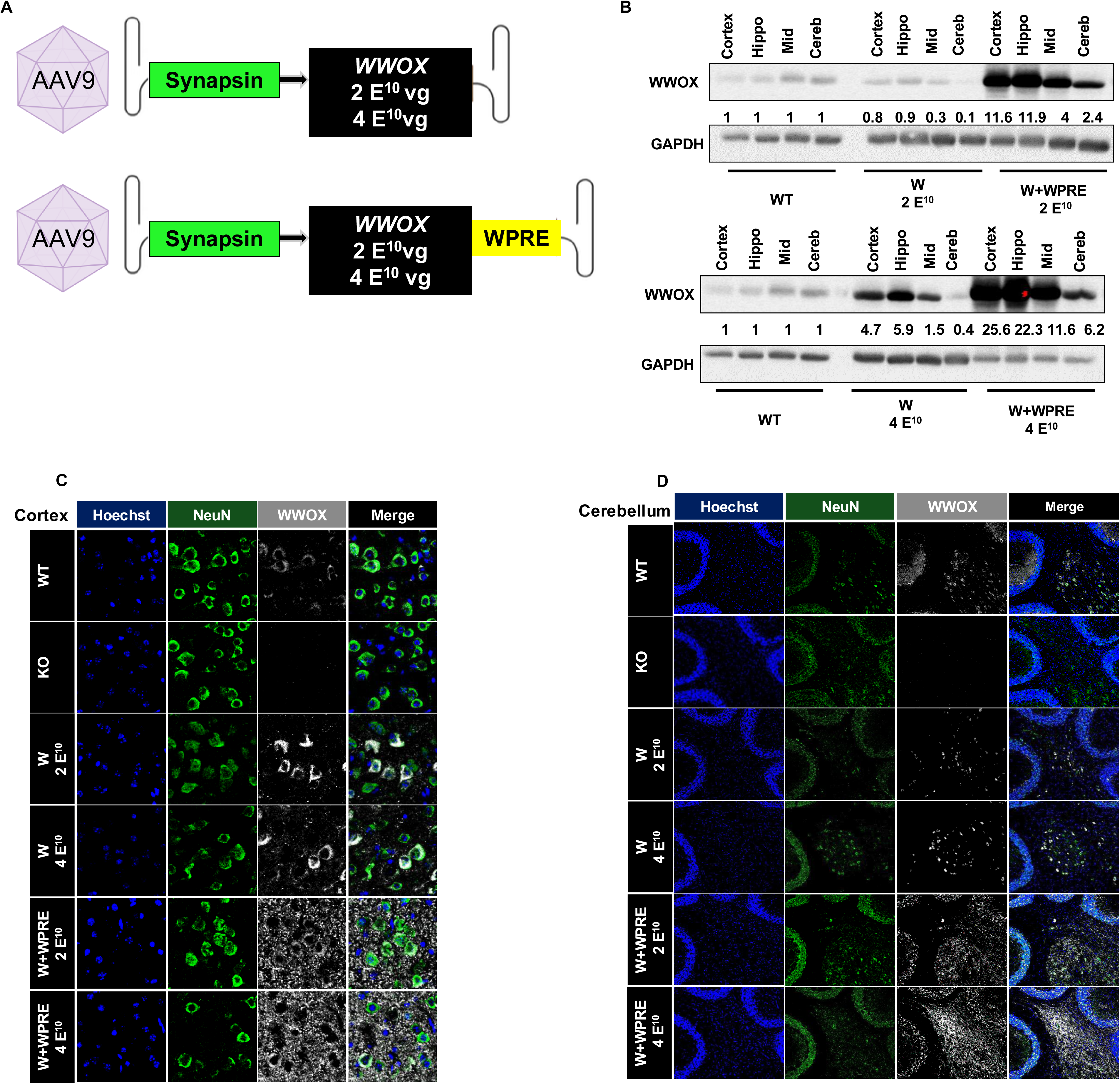
Removal of the WPRE element significantly reduces WWOX overexpression. **(A)** Schematic representation of AAV9 vectors expressing WWOX under the human Synapsin (hSynI) promoter, with or without the WPRE regulatory element. Two viral doses were tested: 2×10^10^vg and 4×10^10^vg. **(B)** Western blot analysis of WWOX protein levels in cortex, hippocampus, midbrain, and cerebellum of WT and KO mice treated with AAV9-hSynI-hWWOX (W) or AAV9-hSynI-hWWOX-WPRE (W+WPRE) at P0. Quantification indicates WWOX intensity relative to WT. **(C–D)** Representative immunohistochemistry images showing WWOX (gray) and the neuronal marker NeuN (green) in the cortex and cerebellum, respectively, WT, KO, and KO mice treated with AAV9-hSynI-hWWOX (W) or AAV9-hSynI-hWWOX-WPRE (W+WPRE). Hoechst (blue) labels nuclei. Scale bars: 50 µm.

To directly assess the impact of WPRE on WWOX levels, we compared AAV-SynI-mediated WWOX expression with or without WPRE following intracerebroventricular delivery at two doses (2×10^10^ vg and 4×10^10^ vg; Figure 2A). Immunoblot and immunohistochemistry analyses revealed that WPRE significantly enhanced WWOX protein levels in a dose-dependent and regionally consistent manner across the cortex, hippocampus (Hippo), midbrain (Mid) and cerebellum (Cereb) (Figure 2B-D, Figure S3A). Notably, WWOX expression driven by the WPRE-containing vector appears more intense and diffuse, with a prominent punctate signal throughout neurons, precluding quantification of WWOX-positive neurons, which may reflect higher expression levels rather than a true change in subcellular localization. In contrast, removal of WPRE resulted in a marked reduction in WWOX expression at both doses, and increasing the vector dose in the absence of WPRE failed to recapitulate the expression levels achieved with lower dose containing WPRE (Figure S3C). Importantly, the non-WPRE vector yields a more uniform, predominantly cytoplasmic WWOX pattern that resembles endogenous expression in WT neurons (Figure 2C, D, Figure S3A). Quantification showed a significant increase in WWOX-positive NeuN⁺ neurons, rising from ∼40% to ∼55–60% following treatment (p < 0.05) (Figure S3B). Together, these findings indicate that limiting WWOX expression by removing WPRE yields a more physiologic neuronal expression profile, underscoring the importance of precise transgene regulation for translational gene therapy approaches.

### WWOX gene therapy improves survival and systemic phenotypes in a dose-dependent manner

Building on our optimization studies, we generated AAV9-hSynI-WWOX vectors lacking the WPRE element to achieve tighter control of transgene expression and to proactively optimize the safety margin for clinical translation. For translational relevance, we evaluated two clinically applicable doses: a lower dose (LD, 1.23×10¹¹ vg) and a higher dose (HD, 2.63×10¹¹ vg) (Figure 3A). Survival analysis revealed a clear dose-response relationship: LD treatment modestly, though significantly, extended lifespan relative to untreated *Wwox*-null controls, whereas HD treatment produced a marked and sustained survival benefit (Figure 3B). Most mice receiving the lower dose that succumbed showed reduced WWOX expression by western blot analysis (Figure S5A-D), suggesting that achieving an adequate level of WWOX expression is critical for effective phenotypic rescue.

**Figure 3:**
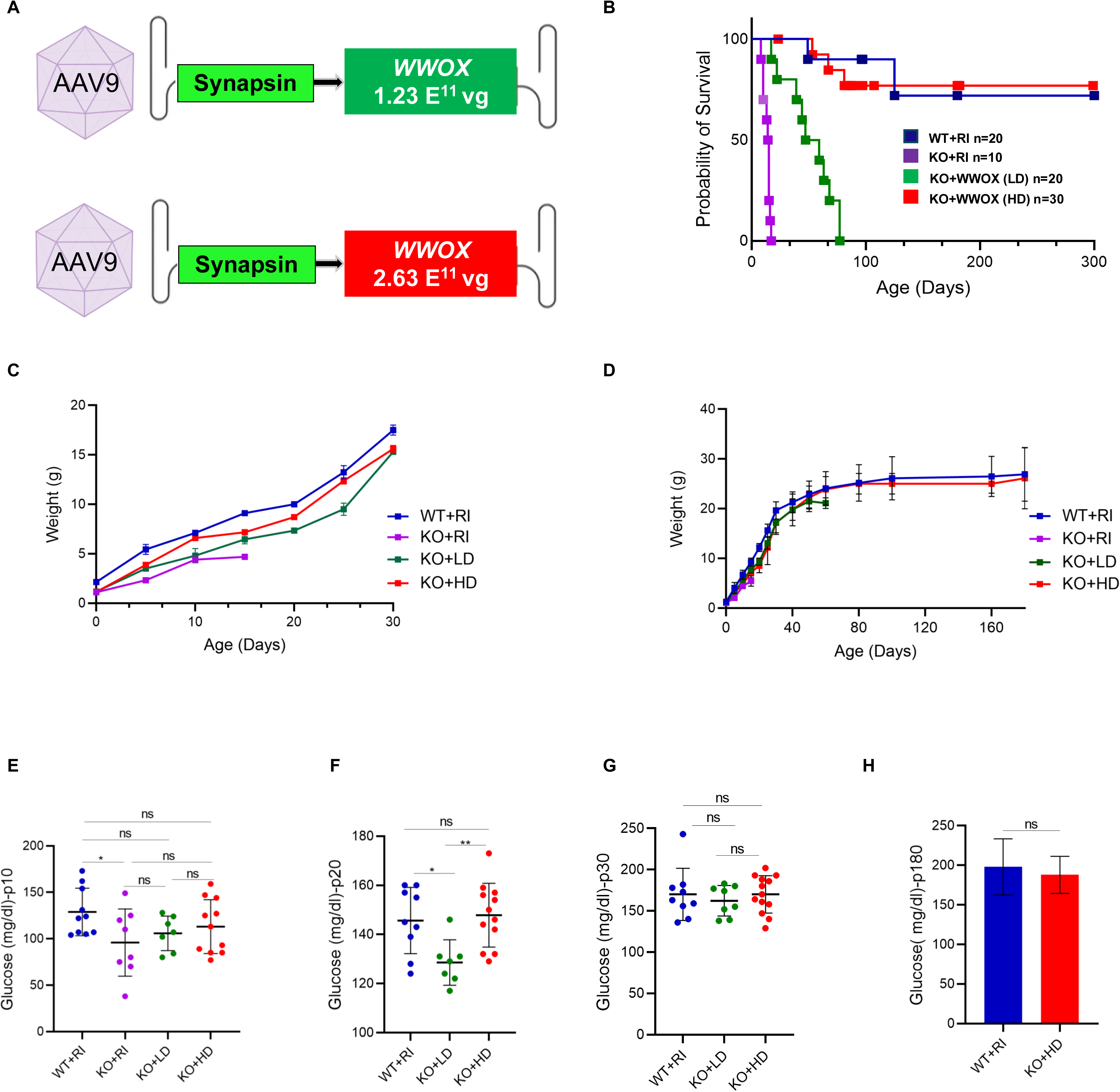
*WWOX* gene therapy enhances survival in a dose-dependent manner. **(A)** Schematic representation of the AAV9-hSynI-hWWOX vectors used for neonatal intracerebroventricular (ICV) injections. Two viral doses were tested: a low dose (LD) (1.23×10^11^vg) and a high dose (HD) (2.63×10^11^vg). **(B)** Kaplan–Meier survival analyses of *Wwox*-null mice treated HD or low-dose LD AAV9-SynI-WWOX compared to KO and WT mice injected with the RI. HD produced survival comparable to WT animals (P=0.78), and significantly extended survival relative to the LD group (P <0.0001) and KO+RI controls (P < 0.0001). LD also significantly improved survival compared with KO+RI controls (P < 0.0001). **(C)** A graph representing body weight during the first postnatal month in KO+LD and KO+HD mice, reaching near WT levels by postnatal day 30, whereas KO mice exhibited marked growth retardation. **(D)** Long-term growth monitoring demonstrated sustained weight gain and normalization in KO+HD mice up to 180 days of age. **(E-G)** Blood glucose levels (mg/dl) measured at postnatal days 10, 20, and 30. **(H**) A Graph representing glucose measurements (mg/dl) at p180 of KO+HD mice were indistinguishable from WT mice. Statistical analysis was performed using Student’s t test (*p<0.05, **p<0.01, ns: not significant), Error Bars represent mean ± SD.

Growth trajectories closely paralleled survival outcomes. Untreated *Wwox*-null mice exhibited severe growth retardation compared with WT controls, while both LD- and HD-treated groups showed progressive weight gain (Figure 3C). By the study endpoint, body weights of HD-treated mice were nearly indistinguishable from WT littermates (Figure 3D), with no significant sex-dependent differences observed.

Given the profound hypoglycemia characteristic of *Wwox*-null mice^45^, we next assessed glucose homeostasis longitudinally. Beginning at postnatal day 10 (P10), both LD and HD treatments partially corrected hypoglycemia. By P20, glucose levels in HD-treated mice were fully normalized and comparable to WT controls, whereas LD-treated mice showed an intermediate improvement. From P30 onward, glucose levels were normalized across all groups, with HD-treated mice maintaining stable WT-like glycemia throughout the study (Figure 3E-H).

Motor coordination was assessed at P18 using hindlimb clasping test ^45^. As shown in (Figures S4A and S4B), untreated *Wwox*-null mice displayed severe motor impairment, whereas LD treatment resulted in partial improvement and HD treatment achieved near-complete rescue, with ataxia scores approaching those of WT controls.

Finally, to determine whether neuronal WWOX restoration ameliorates broader systemic deficits, we assessed reproductive function. Fertility testing using 20 breeding cages per group demonstrated that HD-treated *Wwox*-null mice exhibited fertility rates and litter sizes comparable to WT and heterozygous controls (Figure S4C-E).

Collectively, these data demonstrate that AAV9-hSynI-hWWOX gene therapy confers robust, dose-dependent benefits across survival, growth, glycemic regulation, motor coordination, and fertility. While both doses improved disease phenotypes, the higher dose consistently achieved near-complete rescue, establishing a strong therapeutic window and supporting the clinical potential of precisely tuned neuronal WWOX replacement.

### Neuronal WWOX restoration normalizes neurobehavioral function

To determine whether neuronal WWOX reconstitution rescues behavioral and motor deficits, we assessed *Wwox*-null mice treated with high-dose AAV9-hSynI-WWOX (KO+WWOX HD) and compared them with WT controls (Figure 4A). Behavioral testing could not be performed in untreated *Wwox*-null mice due to severe morbidity and early lethality, and LD-treated mice did not survive to postnatal day 90 (P90); therefore, analyses were limited to WT and HD-treated groups.

**Figure 4:**
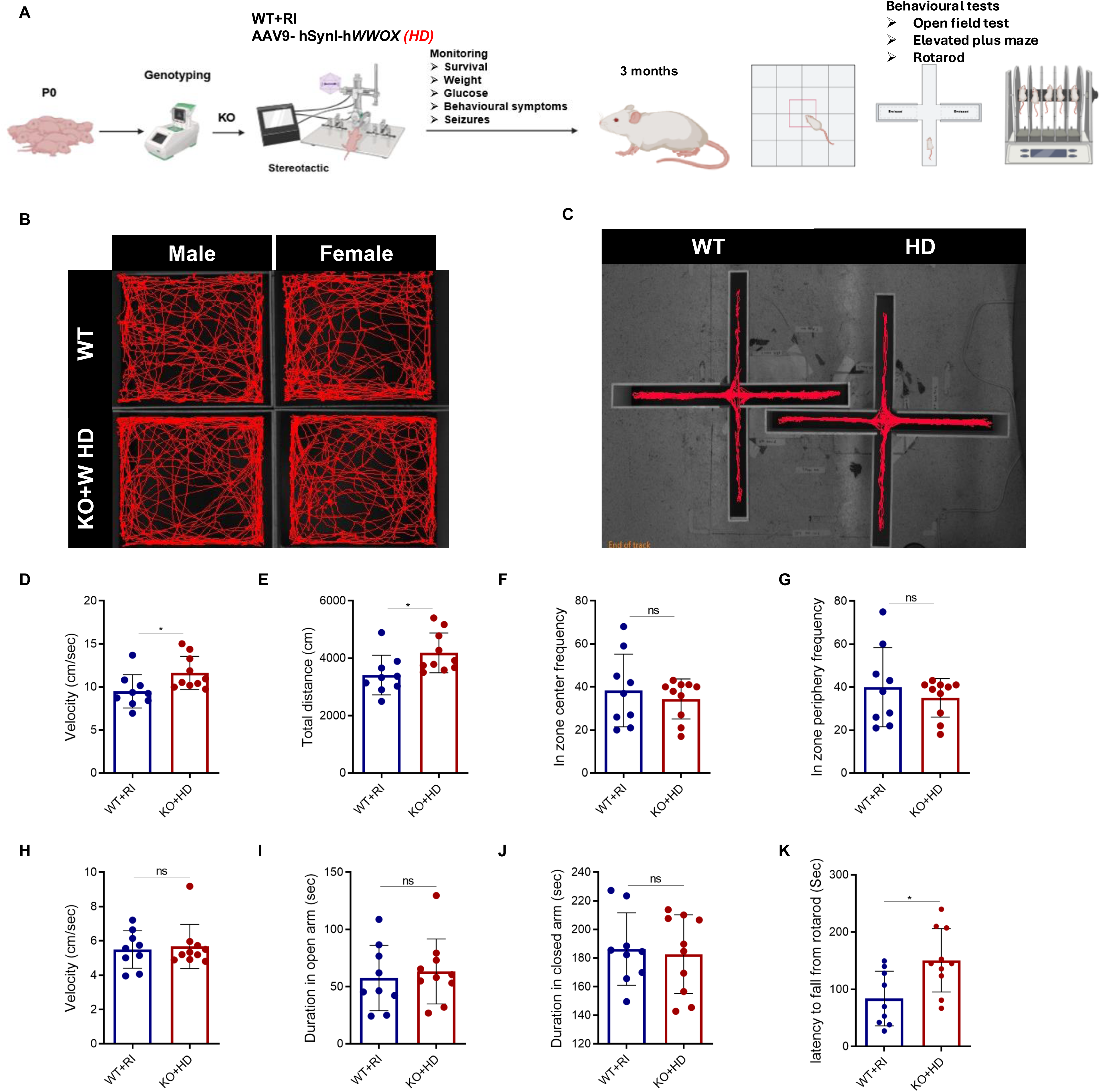
High-dose neuronal WWOX expression restores neurobehavioral performance. **(A)** Schematic representation of the experimental design. KO mice were injected at postnatal day 0-1 (P0-1) with AAV9-hSynI-hWWOX at HD and monitored for survival, body weight, glucose levels, behavioural symptoms, and seizure activity. Behavioural assessments were performed at 3 months of age, including open field, elevated plus maze, and rotarod tests. **(B)** Representative locomotor tracking plots of male and female WT (n=9) and KO+W HD (n=10) mice in the open field test. **(C)** Representative trajectory maps from the elevated plus maze of male and female WT (n=9) and KO+W HD groups (n=10). **(D-G)** Quantitative analysis of open field parameters: velocity (cm/sec), total distance travelled (cm), in zone center and periphery frequency, respectively. **(H-J)** Quantitative analysis of elevated plus maze test parameters: velocity (cm/sec), duration in the open and closed arm (sec), respectively. **(K)** A graph representing the Latency (sec) of the mice (WT=9, KO+W HD=10) to fall from the rotarod. Statistical analysis was performed using Student’s *t*-test (*p<0.05, ns: not significant), Error bars represent mean ± SD.

At three months of age, both male and female HD-treated mice displayed open-field performance indistinguishable from WT controls. Locomotor activity parameters, including movement velocity and total distance traveled, as well as spatial exploration of center and periphery zones, showed no significant differences between groups (Figure 4B, D-G). In the elevated plus maze, HD-treated mice exhibited anxiety-related behaviors comparable to WT mice, indicating normalization of affective responses (Figure 4C, H-J). Motor coordination and learning, assessed by the rotarod test, were indistinguishable between HD-treated mice and WT controls across repeated trials (Figure 4K). Together, these findings indicate that early, high-dose neuronal WWOX gene therapy yielded neurobehavioral and motor outcomes indistinguishable from those of WT controls, encompassing locomotor activity, anxiety-related behavior, and motor coordination.

### AAV9-SynI-WWOX restores WWOX DNA, mRNA, and protein expression in a dose-dependent and durable manner

To assess the efficiency and biodistribution of AAV9-mediated WWOX delivery, we quantified viral DNA (vDNA) levels in brain tissue at postnatal day 30 (P30) using qPCR. Analysis of the cortex, hippocampus, midbrain, and cerebellum revealed a clear dose-dependent pattern, with HD-treated mice exhibiting significantly higher vDNA levels than LD-treated mice across all regions (Figure 5A-D), confirming efficient transduction and dose-dependent vector distribution.

**Figure 5:**
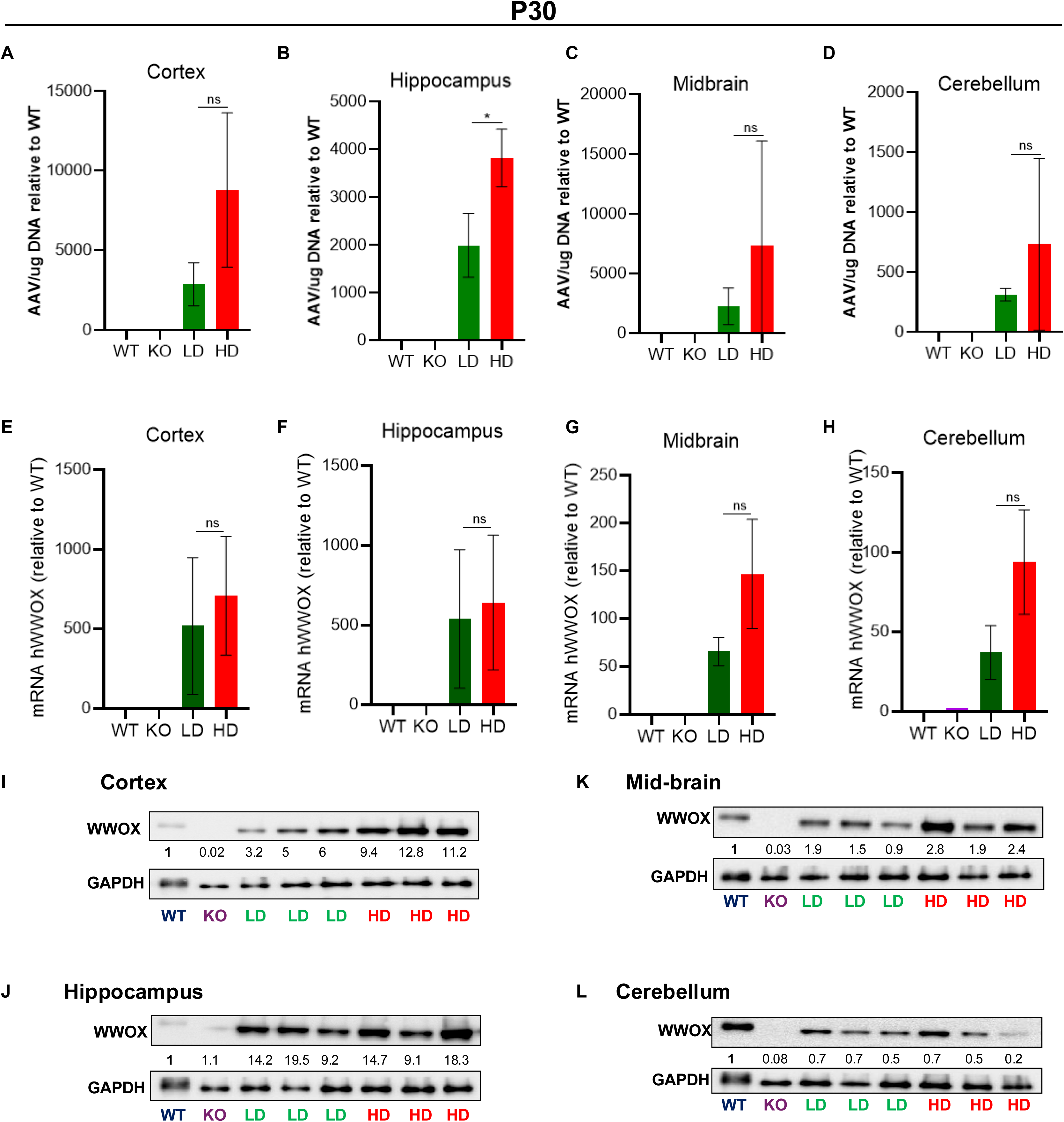
AAV9-hSynI-hWWOX increases WWOX DNA, mRNA, and protein expression in *Wwox*-null mice in a dose-dependent manner. **(A-D)** Vector genome copies (GC) in cortex, hippocampus, midbrain, and cerebellum at postnatal day 30 (P30), measured by qPCR and normalized to WT levels. **(E-H)** WWOX mRNA expression in the same brain regions, normalized to WT. **(I–L)** Representative western blot analysis showing WWOX protein expression in cortex, hippocampus, midbrain, and cerebellum respectively following LD and HD AAV9-hSynI-hWWOX treatment. Quantification indicates WWOX intensity relative to WT. Statistical analysis was performed using Student’s t-test (*p<0.05, ns: not significant, in G, H it is near significant, p-value= 0.08, 0.06 respectively). Error bars represent mean ± SD.

To determine whether the delivered transgene was transcriptionally active, we next measured *WWOX* mRNA levels. RT-qPCR analysis demonstrated robust, region-wide induction of WWOX transcripts in HD-treated mice, whereas LD-treated mice showed lower but readily detectable expression (Figure 5E-H). These data establish that AAV9-hSynI-hWWOX drives efficient, dose-dependent neuronal transcription of the therapeutic transgene.

We then evaluated transgene translation by quantifying WWOX protein levels at multiple postnatal time points. Immunoblot analyses at P30 revealed a marked dose-dependent increase in WWOX protein, most prominently in the cortex and to a lesser extent in the hippocampus, midbrain, and cerebellum, with HD treatment producing substantially higher expression than LD (Figure 5I-L). At later stages (∼3 months), both immunoblot and immunohistochemistry analysis demonstrate that HD-treated mice maintained elevated WWOX protein levels across all examined brain regions (Figure S5A-D, H, I), a pattern that was also evident in the spinal cord (Figure S6A). Notably, mice from either treatment group that failed to survive exhibited reduced WWOX expression, reinforcing the link between effective protein restoration and survival (Figure S5A-D).

Long-term analyses demonstrated sustained WWOX expression. Robust WWOX protein levels persisted at P180 (Figure S5E-G), and widespread, stable expression was maintained at P240 and P300 throughout the cortex, hippocampus, midbrain, cerebellum, and spinal cord following a single neonatal injection (Figure S5J-K, Figure S6B-D). Consistent with physiological expression, WWOX protein was also detected in the sciatic nerve of HD-treated mice (Figure S6E-G), supporting functional relevance in the peripheral nervous system. In contrast, no WWOX expression was detected in the liver following either LD or HD treatment (Figure S6H,I), indicating that transgene expression was restricted to the central and peripheral nervous systems without detectable peripheral leakage.

Together, these results demonstrate that an optimized AAV9-hSynI-WWOX gene therapy enables robust, dose-dependent, neuron-restricted, and sustained restoration of WWOX DNA, mRNA, and protein expression across the nervous system, supporting its durable therapeutic efficacy.

### Loss of WWOX disrupts myelination, while neuronal WWOX restoration promotes dose-dependent myelination and suppresses neuroinflammation

Efficient neuronal signaling depends on proper myelination, a postnatal process that is tightly coupled to neuronal activity. Accumulating evidence indicates that excitatory neurons regulate oligodendrocyte precursor cell (OPC) proliferation and myelin biosynthesis through activity-dependent, non-cell autonomous mechanisms, although the genetic mediators of neuron-oligodendrocyte communication remain incompletely defined^54–66^. Recent studies have implicated WWOX in myelin-related pathologies, including Alzheimer’s disease and multiple sclerosis, where its loss exacerbates neurodegeneration and demyelination^24,25,31^.

To assess the impact of WWOX deficiency and neuronal WWOX restoration on myelination, we performed MBP immunostaining on coronal brain sections from WT, *Wwox*-null, and AAV9-hSynI-hWWOX-treated mice. Consistent with this emerging link, *Wwox*-null mice exhibited profound hypomyelination accompanied by impaired neuronal activity across multiple brain regions, including the corpus callosum (CC), cortex, striatum, anterior commissure (AC) (Figure 6A-E). Strikingly, neuron-specific re-expression of WWOX using AAV9-hSynI-hWWOX resulted in a robust, dose-dependent restoration of myelination. HD treatment achieved near-complete rescue across affected regions, whereas LD treatment produced only partial recovery relative to HD (Figure 6F, Figure S7 I). These data suggest neuronal WWOX as a critical upstream regulator of postnatal myelin development and maintenance.

**Figure 6:**
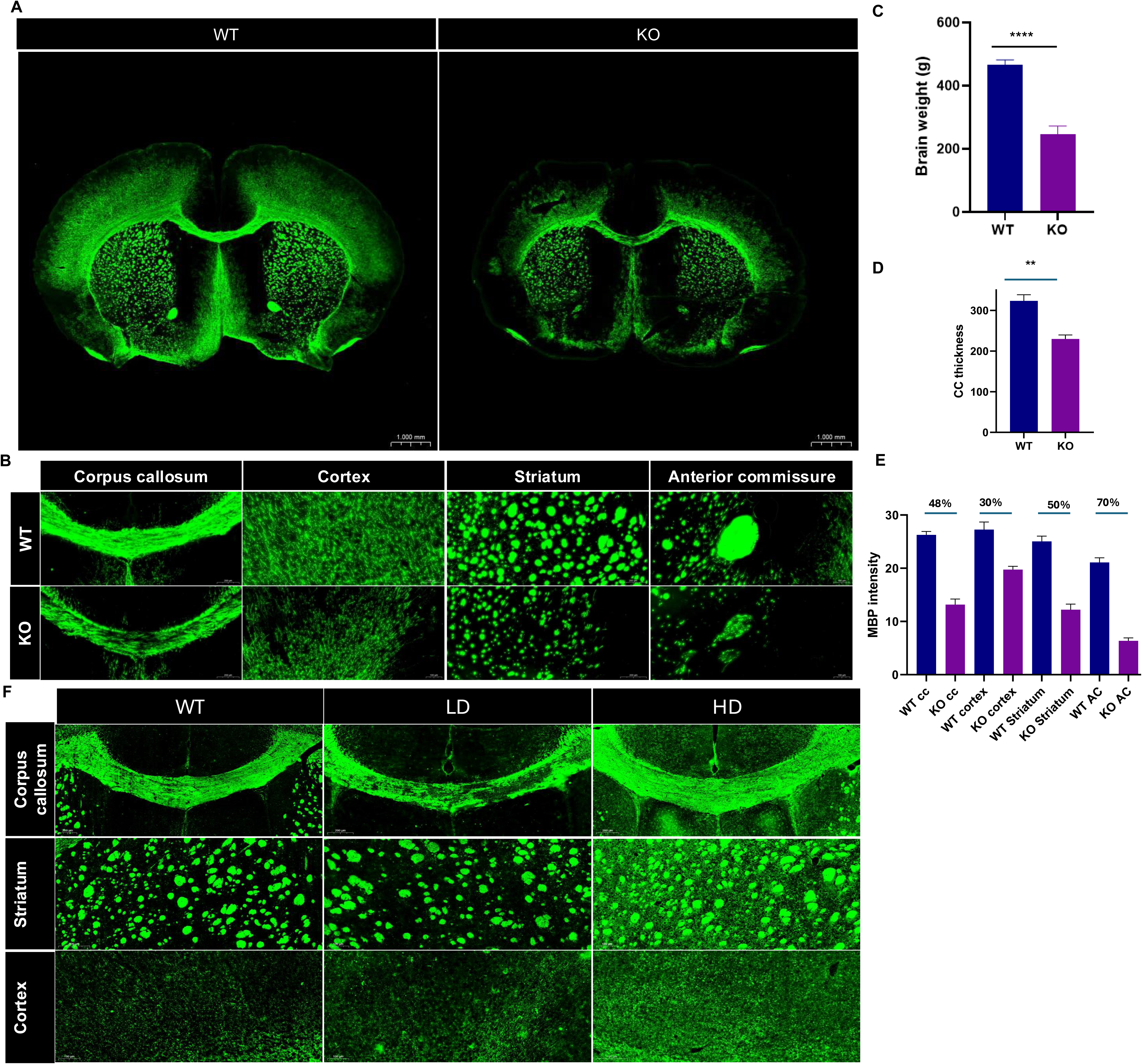
Neuronal WWOX restoration rescues hypomyelination in *Wwox*-null mice in a dose-dependent manner. **(A)** Representative coronal brain sections stained for myelin basic protein (MBP) showing reduced myelination in KO mice compared with WT controls at P18. **(B)** MBP immunohistochemistry staining in the corpus callosum (CC), cortex, striatum, and anterior commissure (AC) of WT and KO mice p18. **(C-E)** Bar graphs showing significant reduction in brain weight (C), CC thickness (D), and MBP intensity across regions (E) in KO mice relative to WT controls. Percentages (E) indicate relative MBP intensity compared to WT. **(F)** Representative MBP staining in WT, AAV9-hSynI-hWWOX-treated KO mice LD and HD, demonstrating dose-dependent restoration of myelination across Corpus callosum (CC), striatum, and cortex. Statistical analysis was performed using Student’s *t*-test (**p<0.01,***p<0.001). Error bars represent mean ± SD.

In parallel, *Wwox*-null mice displayed pronounced astrogliosis and microglial activation, consistent with a heightened neuroinflammatory state (Figure S7A-D). Neuronal WWOX restoration significantly reduced both astrocyte reactivity and microglial density in a dose-responsive manner, indicating effective resolution of neuroinflammation (Figure S7E-H).

Together, these findings demonstrate that WWOX loss disrupts myelin markers and induces gliosis, and neuron-targeted restoration normalizes MBP and glial reactivity in a dose-dependent manner. This dual effect underscores the therapeutic potential of neuronal WWOX gene therapy for WWOX-related neurodevelopmental and neurodegenerative disorders.

### WWOX deficiency promotes early epileptiform activity, while optimized AAV-mediated WWOX delivery suppresses spike-wave discharges

Based on our previous studies demonstrating that both *Wwox*-null and neuron-specific Wwox knockout mice exhibit pronounced epileptiform activity^43^, we sought to directly assess whether HD WWOX gene therapy modulates early network hyperexcitability. To this end, we implemented continuous electrocorticography (ECoG) recordings to quantify epileptiform activity during early postnatal development. Continuous ECoG monitoring was performed in WT and *Wwox*-null pups beginning at approximately postnatal day 14 after ICV administration at P0 of RI or AAV9-hSynI-hWWOX, respectively. Representative ECoG traces from KO pups revealed frequent interictal-like discharges and spike events, whereas WT pups displayed relatively stable baseline activity with sparse spiking (Figure 7A,B). Quantitative analysis of daily spike rates over a 7-day recording period demonstrated a consistently higher incidence of epileptiform spikes in KO pups compared with WT controls (Figure 7B). Averaged spike counts further confirmed a significant elevation in spike activity in KO animals (Figure 7C), indicating that WWOX deficiency leads to early-onset neuronal hyperexcitability during postnatal development.

**Figure 7:**
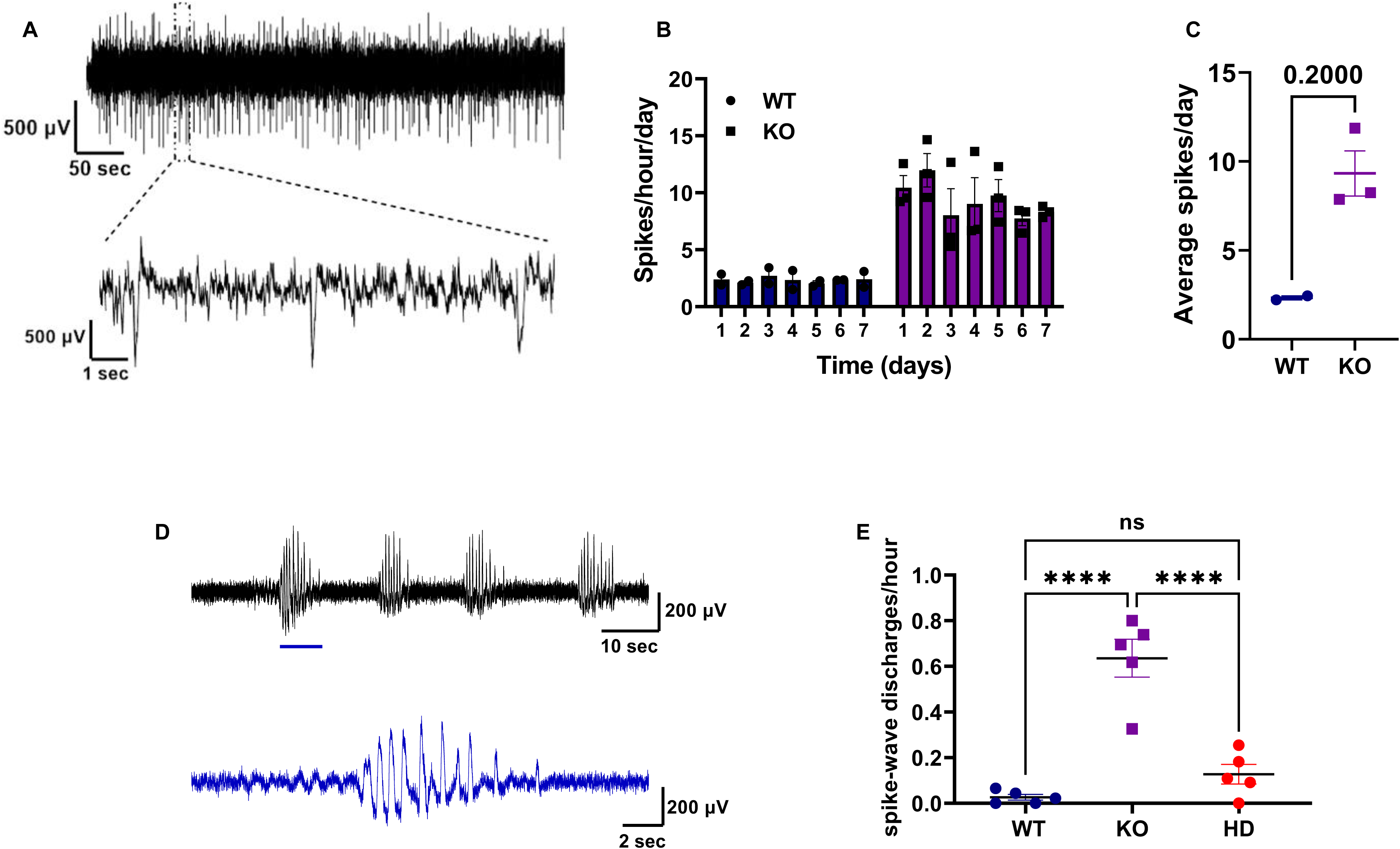
*Wwox* KO littermates exhibit elevated spike activity compared to WT pups, and AAV-mediated WWOX restoration rescues spike-wave discharges (SWDs) in *Wwox* KO pups. **(A)** Representative ECoG traces from WT and WWOX KO pups at ∼P14 showing recurrent spikes. A zoomed-in segment highlights a single spike discharge. **(B)** Quantification of daily spike frequency (spikes/hour) over 7 days of recording. KO pups consistently displayed higher spike rates compared to WT littermates. **(C)** Averaged spike counts across the recording period in WT and KO pups. Data are shown as mean ± SEM; n = 5 littermates per group. Statistical significance was determined using Student’s *t*-test. *p < 0.05; ns, non-significant. **(D)** Representative ECoG trace from a WWOX KO pup at ∼P14 showing a characteristic burst of generalized spike–wave discharges (SWDs). Magnified segment highlights rhythmic SWD activity typical of the KO group. **(E)** Quantification of SWD events per animal across groups (WT, KO, and KO + AAV9-hSynI-hWWOX (HD)). WWOX KO pups exhibited a significantly higher number of SWDs relative to WT, whereas WWOX-rescued pups (HD) showed a marked reduction in SWD incidence. Data are presented as mean ± SEM (n = 5 pups per group). Statistical significance was determined using Student’s *t*-test. ****p < 0.0001; ns: non-significant.

We next assessed whether WWOX restoration could modulate this epileptiform phenotype. In addition to elevated spike activity, KO pups exhibited prominent spike-wave discharges (SWDs), characterized by rhythmic, high-amplitude spike-wave complexes, which were largely absent in WT animals (Figure 7D). Notably, neuronal rescue of WWOX in KO pups using HD AAV-mediated gene delivery markedly reduced SWD incidence, restoring network activity toward WT levels (Figure 7E).

Together, these analyses demonstrate that WWOX loss increases early developmental epileptiform activity, whereas AAV-mediated WWOX restoration effectively suppresses pathological SWDs. Although based on a limited cohort, these findings support a direct role for WWOX in regulating early neuronal network excitability and highlight the therapeutic potential of WWOX gene therapy in mitigating epilepsy-related phenotypes.

### Early postnatal WWOX gene therapy achieves durable therapeutic rescue

All preceding treatments were administered at P0–1. To determine whether later ICV delivery could still confer therapeutic benefit, we performed ICV injection of AAV9-hSynI-hWWOX in *Wwox*-null mice at daily intervals from P0 to P5, with at least three littermates treated per time point. WT littermates received identical injections to control for procedural effects. This developmental period corresponds to rapid brain growth and circuit formation in mice and requires age-specific stereotaxic targeting.

Strikingly, neuronal WWOX restoration using the high-dose vector at any time point between P0 and P5 was sufficient to fully rescue the *Wwox*-null phenotype (Figure S8). Treated mice exhibited a dramatic extension in survival, from approximately three weeks to nearly one year, along with normalization of body weight and blood glucose levels (Figure S8A-D). Moreover, mice showed improved motor coordination and restored fertility (data not shown). Consistently, immunoblot and immunohistochemistry analyses confirmed robust WWOX expression with widespread distribution across major brain regions, including the cortex, hippocampus, and cerebellum (Figure S8E,F). Early WWOX re-expression also markedly enhanced myelination and reduced astrogliosis (Figure S8G-I).

Collectively, these results demonstrate that timely neuronal re-expression of WWOX during early postnatal development is sufficient to reverse the otherwise lethal neurological and glycemia defects of *Wwox* deficiency. Therapeutic rescue beyond this early postnatal window was not explored, as *Wwox*-null mice rapidly deteriorate with progressive neurological dysfunction and early lethality, precluding effective intervention at later stages.

## Discussion

In this study, we establish that optimized AAV9-mediated neuronal replacement of WWOX provides a robust and translationally relevant strategy that rescues the full spectrum of neurological and systemic abnormalities in *Wwox*-null mice. By systematically interrogating promoter choice, cell-type specificity, expression strength, dose, and timing, we define the key parameters governing therapeutic efficacy. Our results establish that neuronal WWOX expression is both necessary and sufficient for survival and CNS homeostasis, whereas expression in oligodendrocytes alone is insufficient to confer benefit.

Gene therapy has demonstrated meaningful clinical success, fundamentally transforming the outlook for patients with previously untreatable genetic disorders. A growing body of work supports AAV-mediated gene replacement as a viable therapeutic strategy for monogenic neurodevelopmental disorders when appropriate cellular targeting and expression control are achieved. Neuron-directed AAV delivery has been shown to rescue structural, electrophysiological, and behavioral phenotypes in mouse models of SCN1A^67^ and SCN1B^68^-associated Dravet syndrome, SLC6A1 deficiency^69^, STXBP1-related disorders^70^, and FOXG1 syndrome^71^, including correction of severe neuroanatomical defects such as corpus callosum agenesis. In parallel, multicenter preclinical studies of MECP2 gene therapy in Rett syndrome have demonstrated robust efficacy across models and favorable safety profiles in nonhuman primates^72^. Collectively, these advances provide compelling evidence that targeted gene delivery to the nervous system can achieve durable gene expression, modify underlying disease biology, and produce clinically meaningful benefits, particularly when intervention occurs early during critical windows of brain development.

In our study, under neonatal AAV9 intracerebroventricular delivery conditions, neuronal WWOX expression driven by the human Synapsin I promoter produced the most robust and sustained therapeutic benefit, whereas ubiquitous or oligodendrocyte-restricted expression resulted in limited or transient rescue. Strong ubiquitous promoters (CMV and CBA) supported early postnatal improvement in survival and glycemic control, indicating that high-level early expression can transiently ameliorate disease manifestations; however, these effects were not maintained at later stages. In contrast, MBP-driven expression failed to confer measurable benefit, which may reflect the limited oligodendrocyte tropism of AAV9 following neonatal ICV administration rather than a lack of relevance for oligodendrocyte WWOX expression. We also cannot exclude that the optimal dose was not achieved, as all vectors were tested at the same titer (4E10). Notably, WWOX expression levels were not normalized across promoter conditions, and vector genome copy number, transcript abundance, or protein levels were not systematically quantified across all constructs. Thus, differences in therapeutic durability could reflect a combination of cell-type specificity, expression magnitude, and temporal kinetics. Nonetheless, the observed promoter-dependent outcomes are consistent with our prior conditional knockout studies, in which deletion of *Wwox* in neural stem/progenitor cells (Nestin-Cre) or postmitotic neurons (Synapsin I-Cre) recapitulated the severe neurological and metabolic phenotypes of global knockout mice, whereas astrocyte- (GFAP-Cre) or oligodendrocyte-specific (Olig2-Cre) deletion produced no overt abnormalities^42^. Taken together, under matched neonatal AAV9-ICV conditions, neuron-restricted WWOX expression emerged as the most effective and durable strategy, while ubiquitous and oligodendrocyte-restricted approaches did not confer sustained rescue within the parameters tested.

Another central insight from the present study is the importance of precise control over transgene expression for both efficacy and safety. Our earlier therapeutic construct incorporated the WPRE element, which is widely used to enhance transgene expression and vector yield. Consistent with this function, inclusion of WPRE markedly increased WWOX protein levels in a dose-dependent and regionally consistent manner across multiple brain areas. However, WPRE-containing vectors also produced a more intense and spatially diffuse neuronal signal, characterized by prominent punctate staining throughout neurons, likely reflecting elevated expression levels rather than altered subcellular targeting. Such amplification may increase peak transgene exposure and introduce variability across developmental stages and brain regions. Although no overt toxicity was observed in prior studies, we removed WPRE as a proactive risk-mitigation step to improve the predictability and physiological fidelity of neuronal WWOX expression for clinical translation. In the absence of WPRE, WWOX expression was reduced but exhibited a more uniform, predominantly cytoplasmic distribution that closely resembled endogenous WWOX patterns in wild-type neurons. We subsequently compensated for lower expression through dose optimization, thereby preserving therapeutic efficacy while maintaining a tighter and more controlled expression envelope.

Using this optimized vector configuration, we identified a clear dose-response relationship, providing strong mechanistic evidence of on-target activity and enabling definition of an optimal therapeutic dose. Neuron-specific WWOX expression driven by the Synapsin I promoter resulted in graded improvement across seizures, ataxia, hypoglycemia, infertility, hypomyelination, neuroinflammation, and premature mortality, with optimal dosing achieving near-complete phenotypic normalization even in the absence of WPRE. Importantly, electrophysiological analyses revealed that *Wwox*-null pups display early-onset neuronal hyperexcitability, characterized by frequent interictal-like spikes and spike-wave discharges, hallmarks of epileptic encephalopathy. AAV-mediated neuronal restoration of WWOX robustly suppressed these pathological discharges, supporting a direct role for neuronal WWOX in maintaining network stability during early brain development (Figure 7). Although conducted in a limited cohort, these findings support the capacity of WWOX gene therapy to correct epileptiform activity at its developmental onset.

An additional key finding is the identification of an early postnatal period during which WWOX restoration is tractable in this murine model. Consistent with experience from other gene therapy approaches targeting severe neurodevelopmental disorders ^67–72^, neonatal or very early postnatal intervention was required to achieve durable rescue. Delivery of AAV9-hSynI-hWWOX within the P0-P5 window normalized growth, markedly extended survival to nearly one year, restored motor and glycemic function, and prevented progressive neurodegeneration (Figure S8). Therapeutic intervention beyond this period was not feasible, as *Wwox*-null mice rapidly deteriorate after P5, developing severe growth retardation, progressive weight loss, and frequent seizures^42,46^. These features rendered the animals highly vulnerable and raised ethical and practical concerns regarding invasive procedures such as intracerebroventricular injection at later stages. Accordingly, the inability to assess later intervention likely reflects a combination of model-specific biological constraints and technical limitations, rather than a definitive boundary for therapeutic responsiveness. These findings should therefore be interpreted in the context of murine neurodevelopment, where rapid postnatal brain maturation and accelerated disease progression may impose a narrower therapeutic window than in humans. In contrast, human patients with WWOX-related encephalopathies may exhibit different developmental trajectories, disease kinetics, and therapeutic responsiveness, potentially allowing for later intervention. Accordingly, future studies should explore alternative strategies to extend the therapeutic window, including earlier prenatal delivery, less invasive or systemic administration routes, combinatorial approaches to stabilize neuronal networks, or adjunct therapies aimed at mitigating seizures, inflammation, or metabolic stress prior to gene replacement. Together, these considerations highlight both the strengths and limitations of the murine model and underscore the need for continued optimization to translate WWOX gene therapy effectively to human patients.

In summary, our study defines a comprehensive therapeutic framework for WWOX-related encephalopathies based on neuronal specificity, dose optimization, and developmental timing. Neuron-targeted AAV9-mediated WWOX replacement achieves durable rescue of both neurological and systemic phenotypes in *Wwox*-null mice, establishing a strong preclinical rationale for translation. Given the established safety profile of AAV9 in pediatric applications, the neuronal precision of the Synapsin I promoter, and the feasibility of neonatal intracerebroventricular delivery, this approach represents a viable path toward clinical intervention in infants with WWOX deficiency. Collectively, these findings position WWOX as a central regulator of neuronal homeostasis and identify its timely restoration as a promising therapeutic strategy for otherwise fatal developmental encephalopathies.

## Acknowledgements

We thank all members of the Aqeilan laboratory and Mahzi team for their contributions and technical support. We are especially grateful to individuals affected by WOREE and SCAR12 syndromes and their families for their continued engagement and generosity. Special thanks to Ms. Takwa Jebara (a former student at the Aqeilan lab), Dr. Kineret Inbar, Ms. Naheel Lawabny, Mr. Wajeeh Salaymeh, and Ms. Marah Bakhtan for technical assistance. This work was supported by Mahzi Therapeutics and the WWOX Foundation.

## Authors Contributions

MO, RA, AB, TB, YW and RIA conceived the study. ICV Gene therapy design and implementation were led by MO. MO, RA, SR, BA designed and performed the in vivo experiments, including phenotyping, molecular analyses, histology/immunostaining, imaging, and behavioral testing. ECoG experiments and analysis were performed by SP, TSA. MO, RA, and RIA wrote the initial manuscript. All authors reviewed and edited the final manuscript.

## Supplementary figure legends

**Supplementary Figure 1: Cell-type-specific WWOX expression driven by different promoters following AAV9 delivery. (A)** Immunohistochemistry staining for WWOX (red) and β3-tubulin (Tuj1) (green) in the cerebellum of KO injected mice using AVV9-hWWOX under different promoters, indicating specificity and intensity of WWOX expression. Hoechst shows nuclear staining. **(B)** Immunohistochemistry staining for WWOX (gray) and CC1 (magenta) in the corpus callosum of KO injected mice using AVV9-hWWOX under different promoters indicating specificity and intensity of WWOX expression. Hoechst shows nuclear staining. **(C)** Immunohistochemistry staining for WWOX (red) and β3-tubulin (Tuj1) (green) in the cerebellum of KO injected mice using AVV9-hSynI-hWWOX indicating specificity and intensity of WWOX expression. Hoechst shows nuclear staining. **(D)** Immunohistochemical detection of GFP demonstrating neuron-specific expression in the cortex, hippocampus, and cerebellum following AAV9-hSynI-GFP delivery using stereotaxic injection.

**Supplementary Figure 2: Promoter-dependent efficacy of AAV9-mediated WWOX gene replacement in *Wwox*-null mice and primary neurons. (A)** Kaplan-Meier survival graph indicates prolonged life span of *Wwox*-null mice injected with AVV9-CBA-hWWOX (4×10^10^ vg) (n=4), compared with KO mice injected with RI (n=6); p<0.05, Log-rank (Mantel-Cox) test. **(B)** Graph demonstrating the body weight (g) of WT (n=6), KO injected with RI (n=6) and KO injected with hWWOX-CBA (n=4) at different postnatal days. **(C)** Bar graphs representing glucose levels (mg/dl) at p14 in WT (n=6), KO injected with RI (n=6) and KO injected with hWWOX-CBA (n=4), p<0.001 compared to KO injected with RI. **(D)** Physical appearance of WT and KO mice injected with AAV9-CMV-hWWOX, AAV9-EF1a-hWWOX, AAV9-MBP-hWWOX and AAV9-hSynI-hWWOX. **(E)** Immunofluorescence staining of β3-tubulin (Tuj1-pink) and WWOX (gray) in primary E16.5 neurons infected with AAV9-hSynI-EGFP, AAV9-hSynI-hWWOX, AAV9-CMV-hWWOX, or AAV9-MBP-hWWOX.

**Supplementary Figure 3: Removal of the WPRE element significantly reduces WWOX expression. (A)** Representative immunohistochemistry images of WWOX (gray) and the neuronal marker NeuN (green) in the hippocampus of WT, KO, and KO mice injected with AAV9-hSynI-hWWOX (W) or AAV9-hSynI-hWWOX-WPRE (W+WPRE). Two viral doses were tested: 2×10^10^vg and 4×10^10^vg. Hoechst (blue) marks nuclei. Scale bars: 50 µm. **(B)** Graphs representing the percentage of NeuN and WWOX positive cells in the cortex, hippocampus and cerebellum in KO mice injected with AAV9-hSyn-hWWOX 2 E^10^ and 4 E^10^. Quantification of the WPRE vector was not possible due to its expression pattern. **(C)** Representative immunohistochemistry images of WWOX (red) in the cortex of WT and KO mice injected with AAV9-hSynI-hWWOX (2.63×10^11^vg) or AAV9-hSyn-hWWOX-WPRE (6×10^10^vg).

**Supplementary Figure 4: Neuronal WWOX restoration improves ataxia in a dose-dependent manner and rescues fertility in Wwox-null mutant mice. (A)** A graph representing quantification of hindlimb clasping in WT, KO injected with RI and AAV9-hSynI-WWOX-treated KO mice receiving LD or HD therapy. Statistical analysis was performed using Student’s *t*-test (***p<0.001, ****p < 0.0001, ns: not significant). Error bars represent mean ± SD. **(B)** Representative images from the clasping test at P18 showing improved motor coordination following neuronal WWOX restoration. **(C-D)** Quantification of reproductive performance in WT, heterozygous (HT), and KO mice treated with AAV9-hSyn-hWWOX (HD). Both the average number of litters per female (C) and the number of pups per litter (D) were comparable across groups. Statistical analysis was performed using Student’s *t*-test (ns: not significant); error bars represent mean ± SD. **(E)** Breeding scheme illustrating fertility rescue in KO mice after WWOX restoration.

**Supplementary Figure 5: AAV9-hSynI-hWWOX increases WWOX protein expression in *Wwox*-null mice in a dose and time-dependent manner at later stages. (A-D)** Western blot analysis of WWOX protein levels across brain regions (cortex, midbrain, hippocampus, and cerebellum) at postnatal day 90 (P90) in WT, KO, and AAV9-hSyn-hWWOX-treated KO mice receiving LD or HD. KO+W mice that did not survive to this age are indicated as “Dead.” Quantification indicates WWOX intensity relative to WT. **(E-G)** Western blot analysis of WWOX protein levels across brain regions (cortex, midbrain and cerebellum) at P180 in WT, untreatedK KO, and AAV9-hSyn-hWWOX-treated KO mice receiving HD. Quantification indicates WWOX intensity relative to WT. **(H)** immunohistochemistry staining of cerebral sections from P90 WT and AAV9-hSyn-hWWOX-treated KO mice receiving LD or HD showing co-localization of WWOX (red) with the neuronal marker β3-tubulin (Tuj1) (green). Hoechst (blue) marks nuclei. **(I)** Representative cerebellar sections showing dose-dependent WWOX expression in WT, KO mice receiving LD or HD at P90. (**J)** Western blot analysis of WWOX protein expression at P300 in KO mice receiving HD. **(K)** Quantification of WWOX protein levels across brain regions at P240 and P300 (n=3). Error bars represent mean ± SD.

**Supplementary Figure 6: Long-term and tissue-specific distribution of WWOX expression following ICV AAV9-hSynI-hWWOX delivery. (A)** Western blot analysis of WWOX protein levels from spinal cord extracts (cervical region) of AAV9-hSynI-hWWOX treated KO mice at LD and HD compared to WT littermates. Quantification indicates WWOX intensity relative to WT. **(B)** Immunohistochemistry staining of spinal cord sections from HD-treated mice at P300 showing co-localization of WWOX (red) with the neuronal marker β3-tubulin (Tuj1) (green). Hoechst (blue) marks nuclei. **(C-D)** Western blot analysis of WWOX protein expression at P240 and P300, demonstrating stable protein expression over time. Quantification indicates WWOX intensity relative to WT. **(E-F)** Western blot analysis of peripheral nervous system (PNS) tissue (sciatic nerve), of HD-treated mice compared to WT and KO mice. **(G)** Immunohistochemistry staining of sciatic nerve sections confirming neuronal WWOX expression (colocalization of WWOX (red) with β3-tubuli (Tuj1) (green). **(H-I)** Western blot analysis of liver samples at P90 and P180, respectively. Quantification indicates WWOX intensity relative to WT.

**Supplementary Figure 7: Neuronal WWOX restoration reduces neuroinflammation and astrogliosis in Wwox-null mice and improves myelination in a dose-dependent manner. (A)** Representative immunohistochemistry of Iba1 (magenta) in WT and KO mice at p20. **(B)** Graph represents percentage of Iba1 area in KO compared to WT shown in A. **(C)** Representative immunohistochemistry of GFAP (green) in WT and KO mice at p20. **(D)** Graph represents percentage of GFAP area in KO compared to WT shown in C. **(E-H)** immunohistochemistry staining of Iba1 and GFAP immunoreactivity in WT, AAV9-hSynI-hWWOX-treated KO mice both LD and HD treatment. F and H show differential quantification of microglia (F) and astrocytes (H) in each treatment. **(I)** Representative MBP immunostaining (gray) at ∼P90 in KO+W LD and KO+W HD mice relative to WT-RI. Hoechst (blue) marks nuclei. Statistical analysis was performed using Student’s *t*-test (*p<0.05, ***p<0.001). Error bars represent mean ± SD.

**Supplementary Figure 8. Early postnatal AAV9-mediated neuronal WWOX delivery improves survival and corrects systemic and CNS phenotypes in *Wwox*-null mice. (A)** Kaplan-Meier survival analysis (till P40) of KO mice treated with HD at P1-P5 (n=3-7) compared to KO and WT mice injected with RI (n = 6). **p<0.001, log-rank (Mantel–Cox) test. **(B)** Kaplan-Meier survival analysis (till P300) of KO mice treated with HD at P1-P5 (n=3-7) compared to KO and WT mice injected with RI (n = 6). **p<0.001, log-rank (Mantel–Cox) test. **(C)** Graphs showing body weight (g) of WT (n = 6), KO mice injected with RI (n = 6), and KO mice injected with HD at P1-P5 (n=3-7) at postnatal day 14 (P14). **(D)** Bar graph representing glucose levels (mg/dl) at p14 in WT (n=6), KO injected with RI (n=6) and KO injected HD at P1-P5 (n=3-7). **(E)** Representative images of immunostaining for WWOX (red) and the neuronal marker β3-tubulin (Tuj1) (green) from KO mice treated with HD at P5 and collected at p30 for analysis. Hoechst (blue) marks nuclei. **(F)** Western blot analysis of WWOX protein expression in the cortex of KO mice treated with HD at P5 and collected at p30 for analysis. **(G)** Representative images of MBP immunostaining from KO mice treated with HD at P5 and collected at p30 for analysis showing improved myelination in the cortex, hippocampus, cerebellum, striatum, and nucleus accumbens compared to KO mice. **(H)** Representative GFAP immunostaining showing elevated astrocyte reactivity in KO brains, which was normalized in KO mice injected at P5. **(I)** Quantification of GFAP-positive from the images in H. Statistical analysis was performed using Student’s *t*-test (**p<0.01, ***p<0.001, ns: not significant). Error bars represent mean ± SD.

